# Testicular sex cord–stromal tumors in mice with constitutive activation of PI3K and loss of Pten

**DOI:** 10.1101/2024.09.16.613175

**Authors:** Marija Dinevska, Lachlan Mcaloney, Samuel S. Widodo, Gulay Filiz, Jeremy Anderson, Sebastian Dworkin, Simon P. Windley, Dagmar Wilhelm, Theo Mantamadiotis

## Abstract

Testicular tumors are the most common malignancy of young men and tumors affecting the testis are caused by somatic mutations in germ or germ-like cells. The PI3K pathway is constitutively activated in about one third of testicular cancers. To investigate the role of the PI3K pathway in transforming stem-like cells in the testis, we investigated tumors derived from mice with post-natal, constitutive activation of PI3K signaling and homozygous deletion of tumor suppressor *Pten*, targeted to nestin expressing cells. Mice developed aggressive tumors, exhibiting heterogeneous histopathology and hemorrhaging. The tumors resemble the rare testis tumor type, testicular sex cord–stromal Leydig cell tumors. Single cell resolution spatial tissue analysis demonstrated that T-cells are the dominant tumor infiltrating immune cell type, with very few infiltrating macrophages observed in the tumor tissue, with CD8^+^ T-cells predominating. Further analysis showed that immune cells preferentially localize to or accumulate within stromal regions.

**Graphical Abstract:** 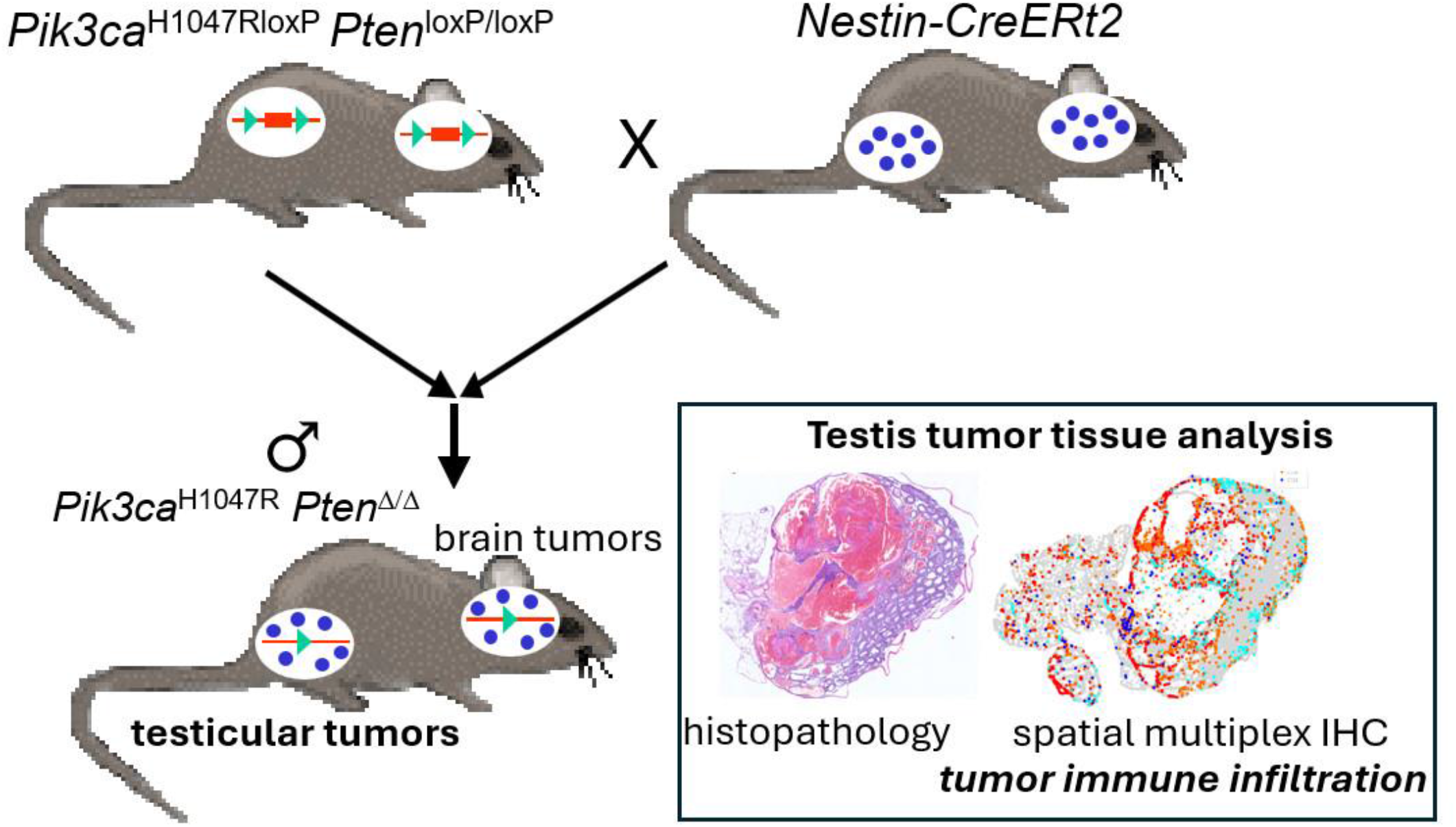

## 1 Introduction

Testicular cancers are the most common malignancy of young men and the most common cancer in young men of European ancestry, where of the 71,000 new cases in 2018, more than one-third were recorded in Europe (1)(2). Despite the improvement in patient survival, patients can experience serious complications due to the effects of chemotherapy, radiation, and surgery, including sterility. Testicular cancer is thought to arise from several progenitor cell types, depending on the type of testicular cancer. Germ cell tumors are the most common type of testicular tumor, with sex cord stromal tumors representing a rarer form of testicular cancer. Within each of these types of tumors, lie further subtypes that are characterized by different cellular and histopathological features. Of the sex cord stromal tumors, Leydig cell cancer is the most common (3). Malignant Leydig cell tumors respond poorly to conventional therapy (4), with a median post-therapy survival time of two years, ranging between a few months to more than ten years, and manifestation of distant metastasis in many cases (3,5).

Genomic analysis shows that *KIT, KRAS* and *NRAS* somatic mutations were associated with seminoma oncogenesis (6). *KIT* encodes a receptor tyrosine kinase and is the receptor for stem cell factor (SCF/KITL) (7). The dominant somatic mutation in KIT wild-type seminomas involves mutation of the *PIK3CA* gene (6). *PIK3CA* encodes for the phosphoinositide 3-kinase (PI3K) catalytic alpha subunit, p110α. The PI3K pathway is one of the major cell signaling activities downstream of KIT activation, and is a major regulator of cell proliferation and cell survival. Almost 30% of testicular tumors harbor somatic mutations in genes linked to the regulation of the PI3K pathway (8). The PI3K pathway negative regulator and tumor suppressor PTEN is implicated in 4% of testicular germ cell tumors, where half of the cases are homozygous deletions and the remaining are a mixture of missense, splice or truncating mutations (9). Aberrant upregulation of the PI3K pathway is a feature of almost all cancer types, given that the pathway is a key regulator of cell survival and proliferation (10).

Despite the identification of somatic mutations associated with testicular germ cell tumors, there is little information on the genetics of Leydig cell tumors, in part due to the rarity of this type of cancer and lack of sufficient tumor tissue samples that are amenable to meaningful genomic analysis. To address the role of the PI3K pathway in testicular cancer, we investigated tumors derived from mice with constitutive activation of PI3K signaling in Nestin expressing cells, generated by the inducible expression of a *Pik3ca*^*H1057R*^ mutation in combination with homozygous deletion of *Pten* (11). These mice were developed with the primary aim of investigating the role of PI3K signaling in neural stem progenitor cell-derived brain tumors, but we discovered that a subset of mice carrying the *Pik3ca* and *Pten* mutations exhibited tumors within the testes. Here, we report the development of aggressive testicular tumors that are derived from interstitial cells, in mice with constitutive PI3K activation and deletion of *Pten*, and show that these tumors exhibit histopathological features of Leydig cell tumors and cytotoxic and helper T-cell infiltration.

## 2 Methods

### 2.1 Ethics statement

Mouse experiments were carried out with the approval of The University of Melbourne, School of Biomedical Sciences (AEC No 1112336.1).

### 2.2 Mice

Mice heterozygous for a latent Cre recombinase (Cre)-inducible knock-in of the *Pik3ca*^*H1047R*^ mutation (*Pik3ca*^*H1047R-loxP*^) (12) and/or two Cre-inducible *Pten* deletion alleles (*Pten*^*loxP/loxP*^) (13) were crossed with mice expressing a single (heterozygous) *Nestin-CreER*^*T2*^ transgene (14). Mice were housed and genotyped, as previously described (11).

### 2.3 Tissue staining and immunostaining

For immunohistochemical analysis, paraffin embedded tissue sections were cut at a thickness of 4 μm, cleared of paraffin and rehydrated through an ethanol to water series. Antigen retrieval was performed by incubating the slides in a citrate buffer (10mM Citric Acid, 0.05% (v/v) Tween 20, pH 6.0) or Tris EDTA pH 9.0 for 20 minutes at 95^°^C. Endogenous peroxidase activity was blocked using 3% Hydrogen peroxide (Sigma-Aldrich, Missouri USA) for 10 minutes at room temperature. Slides were incubated overnight at 4°C with antibodies directed against nestin (Abcam, ab11306), cytochrome P450 cholesterol side chain cleavage enzyme (SCC / Cyp11a1) (15), mouse Vasa homolog gene (Mvh / Ddx4) (Abcam ab270534), Sox9 (16), phospho-Erk1/2 (Cell Signaling Technologies 4370), phospho-Akt (Cell Signaling Technologies 4060) and phospho-Creb (Abcam ab32096). Slides were then washed in phosphate buffered saline with Tween 20 (PBST) and incubated with a biotinylated goat anti-polyvalent universal antibody (Abcam ab93677) for 10 minutes. Slides were then incubated with Streptavidin peroxidase complex (Abcam) for 10 minutes at room temperature before being washed in phosphate buffered saline, 0.5% Tween-20, pH 7.4. Slides were incubated with DAB (3,3’-diaminobenzidine) solution until a color change was observed. Slides were then washed in tap water for 2 minutes, then counterstained using hematoxylin and Scott’s tap water. Slides were passed through increasing concentrations of ethanol (70%, 90% 100%) and xylene, then coverslips overlaid onto DPX mounting media (Sigma-Aldrich).

### 2.4 Multiplex immunohistochemistry

Multiplex IHC was performed using an Opal 7-Color IHC Kit (Akoya Biosciences) on a Bond RX automated stainer (Leica Biosystems). All primary antibodies were diluted in the Opal Blocking/Antibody Diluent. An antibody panel was used to investigate immune cell localization in the tumors. The antibody panel included CD68 (Abcam ab955), pCREB (Abcam ab32096), CD3 (Abcam, ab16669), CD4 (Abcam ab133616), CD8 (Abcam ab4055) and Nestin (Abcam ab11306).

4µm formalin-fixed paraffin-embedded (FFPE) tissue sections were baked at 60°C and deparaffinized in xylene and rehydrated in a serial dilution of ethanol. All slides were then subjected to heat-induced antigen retrieval prior to incubation with 3% hydrogen peroxide. Primary antibody was then added to tissue and incubated at room temperature for 60 minutes, followed by incubation with Opal Polymer HRP Ms + Rb for 30 minutes. Immunofluorescent signals were visualized using Opal fluorophores (Opal 520, 540, 570, 620, 650 or 690), diluted at 1:150 in Plus Automation Amplification Diluent. Serial multiplexing was performed by repeating antigen retrieval, primary and secondary antibody incubation, and Opal polymer visualization. Slides were then counterstained with DAPI and coverslipped using ProLong Glass Antifade Mountant (Thermofisher).

### 2.6 Microscopy and image analysis

Multiplex IHC images were acquired using the Vectra 3.0.5 Multispectral Imaging Platform (Perkin Elmer, USA) at 40x magnification. Spectral deconvolution was then performed using inForm 2.4.4 (Akoya Biosciences). High-resolution multispectral images were then fused and analyzed using the software HALO (Indica Labs). Image analysis was carried out using HALO, where cell phenotyping was performed using the analysis algorithm Highplex FL 3.0.3, according to the cell phenotyping matrix. Cell phenotyping involved adjusting the nuclear detection sensitivity, establishing the minimum intensity threshold for positive detection, and defining biomarker localization as nuclear, cytoplasmic, or membranous.

### 2.7 Statistical analysis

Statistical analyses included a non-parametric One-way ANOVA test and Wilcoxon matched pairs signed rank test with GraphPad Prism v.9 software, as indicated in each figure legend. Statistical significance is represented by ***(p<0.001) and ****(p<0.0001).

## 3 Results

### 3.1 Nestin expressing cells are located in the testicular interstitium

To identify Nestin expressing cells in healthy testis we performed immunohistochemical analysis on mouse tissue and showed that most Nestin-expressing cells were located in the interstitial tissue (Fig. 1A). To identify the major Nestin-expressing cells in human testis, we investigated Human Protein Atlas (17) data, which indicated that interstitial cells also exhibit the strongest Nestin expression (Fig. 1B).

**Figure 1.**
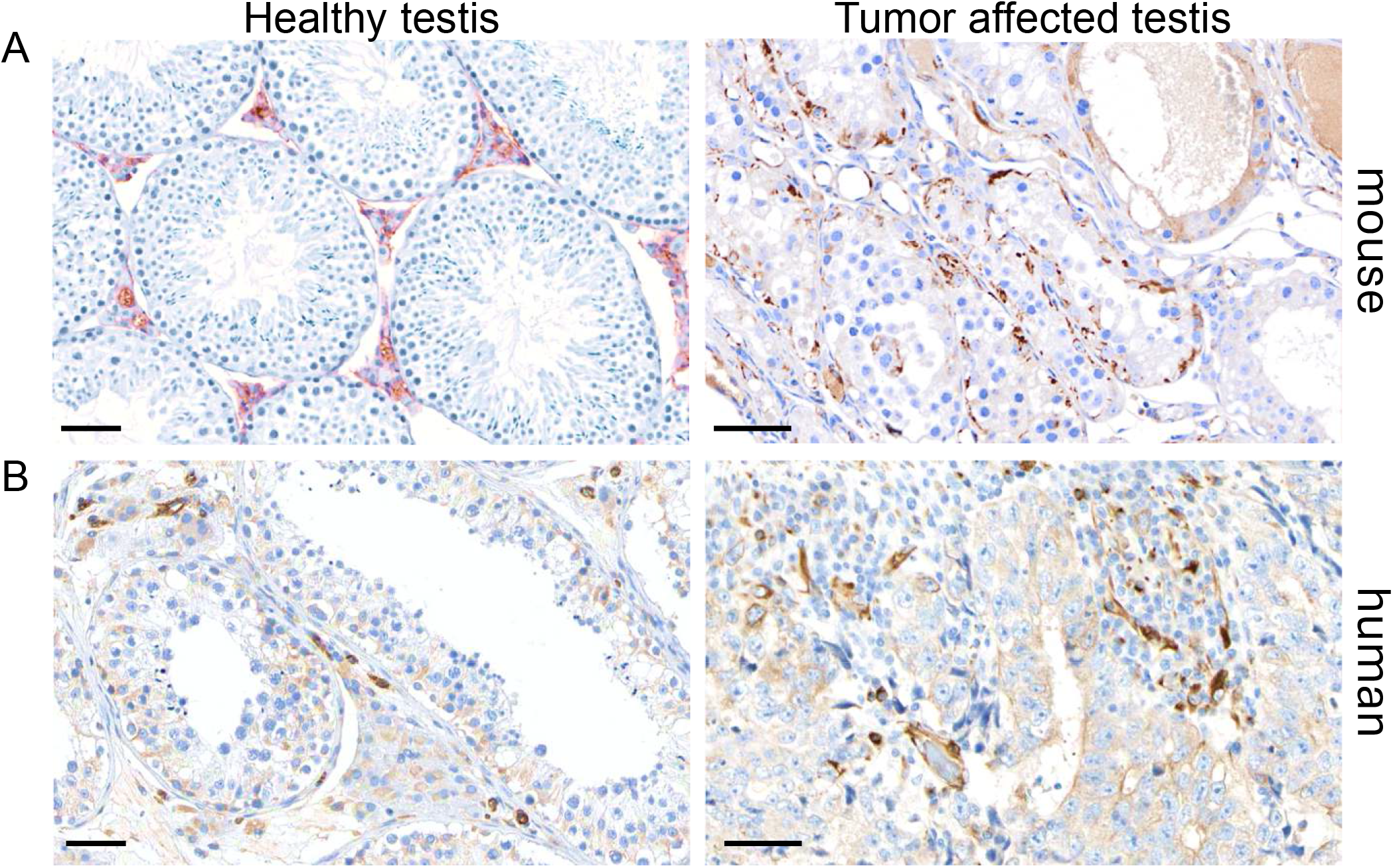
Nestin expression (brown staining) in healthy non-tumor and tumor mouse (A) and human (B) testicular interstitial cells. Hematoxylin (blue staining) was used to counterstain the tissue sections. Tumor tissues exhibit disrupted seminiferous tubule morphology and disrupted germ cell development. Mouse tumor tissue is from the *Pick3ca-Pten* mutant mice (11) and human tissue images are from the Human Protein Atlas version 23.0 *(proteinatlas*.*org)*, testis healthy and tumor tissue panel (17). Scale bars are 50μm.

### 3.2 Constitutive activation of PI3K signaling in testis interstitial cells leads to tumor development

From a total of 49 adult (8 weeks old) male mice carrying a heterozygous or homozygous *Pik3ca*^*H1047R-lox*^ allele, homozygous *Pten*^*loxP/loxP*^ alleles and mono-allelic for the *Nestin-CreER*^*T2*^ transgene, six exhibited overt unilateral testicular enlargement. Upon dissection, gross morphology showed the affected testis was between 1.5 and 2 times the size of the unaffected testis (Fig. 2A). Testes with tumors invariably exhibited regional hemorrhage (Fig. 2B). Histological examination showed that most of the normal cellularity and seminiferous tubules were absent, with some normal seminiferous tubules observed at the periphery, and necrosis with acellular spaces filled with blood (Fig. 2B and C). In tumor affected testes that harbored some intact seminiferous tubules, we observed the presence of scarce, elongated spermatids, instead these tubules which contained sloughed off spermatogenic and Sertoli cells (Fig. 2C).

**Figure 2.**
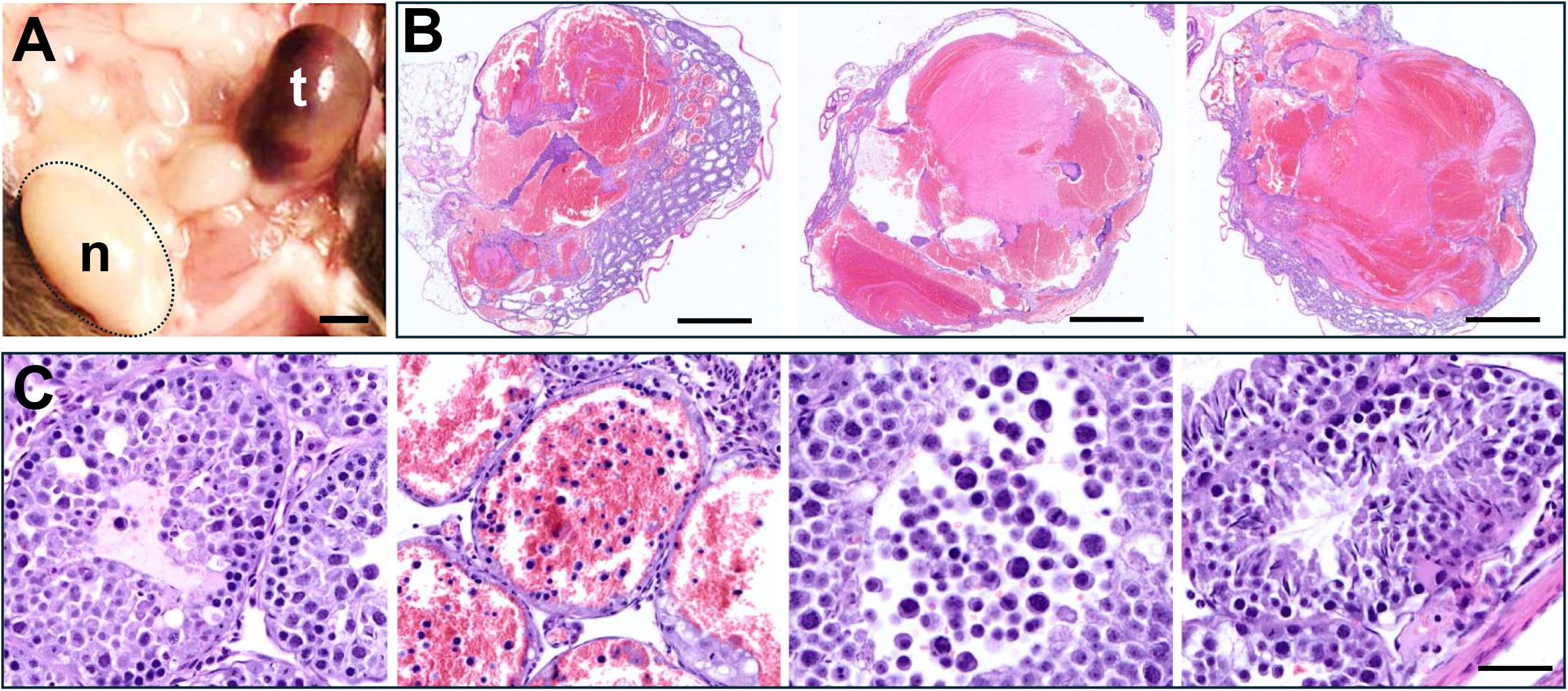
Severe disruption of tissue morphology and hemorrhage in testes with tumors. (A) Dissected mouse showing the testis with a tumor and hemorrhaging (t), and a normal left testis (n). Scale bar is 5mm. (B) Three hematoxylin and eosin-stained testis tissue sections with tumors from different mice. Tumor affected testes exhibit large regions of hemorrhage and stroma, with intact seminiferous tubules evident around the periphery of the tumor tissue. The scale bar is 1mm. (C) Seminiferous tubules from tumor affected testes showing immature germ cells throughout, as well as blood-filled seminiferous tubules with germ cell remnants, immature pleomorphic cells, and cyst-like structures, with abnormally differentiated spermatocytes. Scale bar is 50μm.

### 3.3 Cell type analysis and cell signaling activation in testicular tumors

To identify major testis cell types we used immunohistochemistry (IHC) and antibodies recognizing germ cells (Mvh), Lyedig cells (Cyp11a1) and Sertoli cells (Sox9). Mvh IHC shows that germ cells are present in the few seminiferous tubules in the tumor-affected testis, but germ cell morphology is abnormal, where the germ cells are larger, compared to germ cells in healthy testis (Fig 3). Cyp11a1 IHC was observed in interstitial cells of both tumor-affected and healthy testis. Weak Sox9 staining was seen in the tumor-affected testis, compared to healthy testis.

**Figure 3.**
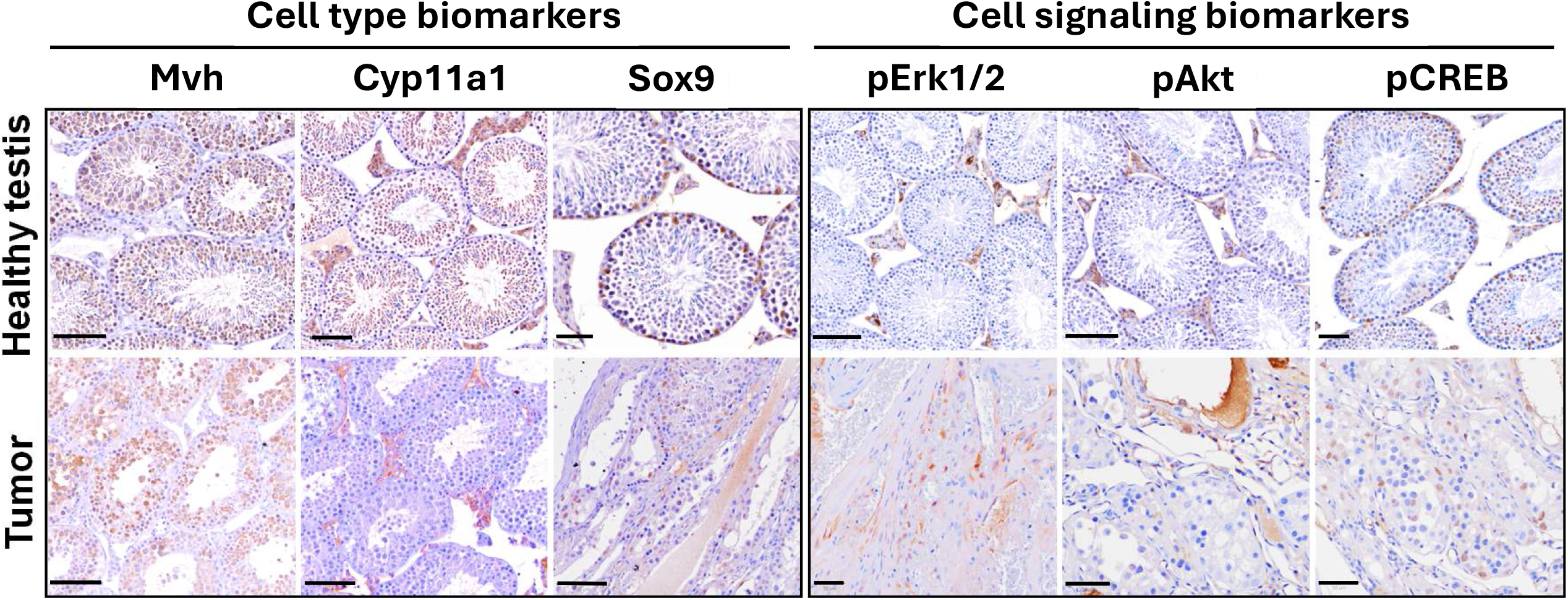
Cell type and cell signaling biomarker expression in healthy non-tumor and *Pik3ca-Pten* mutant tumor testis, identified by immunohistochemistry. Mvh-expressing germ cells, Cyp11a1-expressing Leydig cells, and Sox9-expressing Sertoli cells in healthy mouse testis and tumor affected testis. Activation of PI3K signaling, MAPK signaling and CREB in testis tumors was assessed using phospho-specific antibodies for Akt, Erk (1/2) and CREB, respectively. Scale bars are 50μm, as indicated.

To assess the activation of PI3K and MAPK signaling, which are constitutively upregulated in tumors, we used IHC to assess the phosphorylation status of Akt (PI3K pathway biomarker) and Erk1/2 (MAPK pathway biomarker) (Fig. 3). The expression of phospho-CREB (pCREB) was also measured. CREB, a kinase-inducible transcription factor, is phosphorylated when activated by upstream cell signaling pathways, including cAMP, PI3K and MAPK, and is also used to identify proliferating cells in the testis (18). Upregulated PI3K (pAkt) and MAPK (pErk1/2) activation was evident in both healthy and tumor affected mouse testis tissue, localized within the interstitium (Fig. 3). The expression of pCreb is consistent with the activation of transcriptional programs regulating cell proliferation.

### 3.4 T-cells are the dominant tumor infiltrating immune cell in testicular tumors

Tumor infiltrating immune cells are a feature of many cancers, and with the increasing application of immunotherapy, knowledge of the specific immune cell composition and anatomical location of the immune cells can provide clinically relevant information for tumor- and/or patient-specific immunotherapy. To determine tumor immune cell composition, multiplex immunohistochemistry was performed to identify CD4 and CD8 T-cell subsets and macrophages, and their localization with respect to stromal and tumor cell rich regions (Fig. 4). Spatial analysis showed that tumors harbored more CD8+ T-cells (83% of all immune cells), compared to CD4+ T-cells (17% of all immune cells), and that both T-cell subsets were preferentially localized in tumor-cell rich regions, compared to hypocellular stromal regions. CD8+ T-cells comprised 0.3% of all stromal cells and 2.2% of all cells in the tumor cell-rich regions, and CD4+ T-cells comprised approximately 0.3% of all stromal cells and 2.2% of all cells in the tumor cell-rich regions. Macrophages were rarely observed, with the complete absence of macrophages in many sections examined. Nestin expressing cells and pCREB expressing cells were present in tumor-cell rich regions, comprising 1.8% and 0.7% of all cells (Fig. 4).

**Figure 4.**
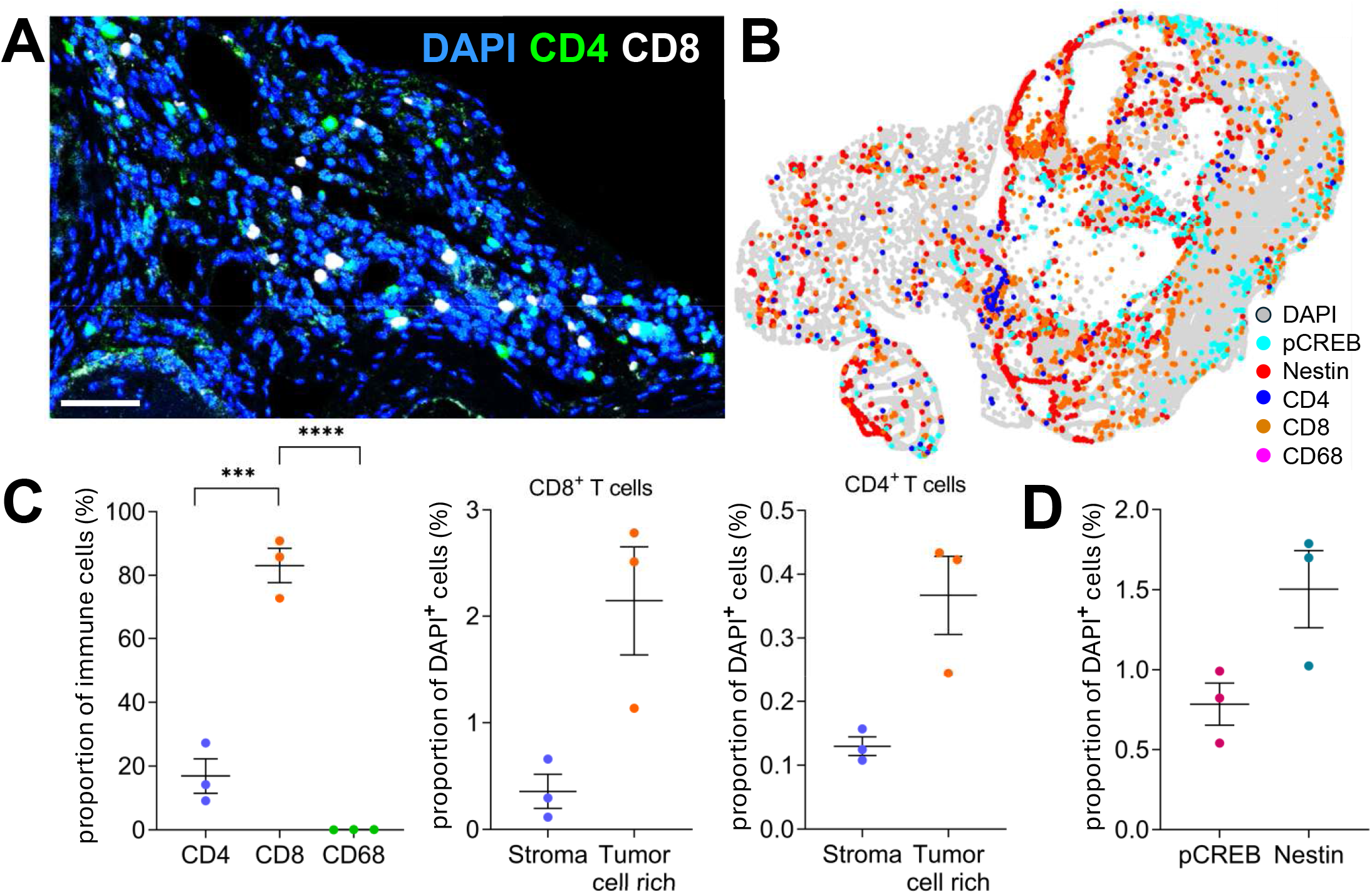
T-cells are the dominant tumor infiltrating immune cell type in the *Pik3ca-Pten* mutant mouse testis. (A) CD4^+^ (green) and CD8^+^ (white) T-cells are localized to tumor cell-rich regions. (B) Spatial relationship of immune cells and tumor cells. CD4^+^ and CD8^+^ T-cell subsets, CD68^+^ macrophages, pCREB and nestin antibodies label proliferating and immature undifferentiated tumor cells, respectively. (C) Quantification of immune and tumor cells. The left panel compares T-cell and macrophage proportions and shows that tumors harbor more CD8^+^ T-cells (83% of all immune cells), compared to CD4^+^ T-cells (17% of all immune cells). The panels on the right show that both T-cell subsets preferentially localize to tumor-cell rich regions, compared to hypocellular stromal regions. CD8^+^ T-cells comprised 0.3% of all stromal cells and 2.2% of all cells in the tumor cell-rich regions, and CD4^+^ T-cells comprised approximately 0.3% of all stromal cells and 2.2% of all cells in the tumor cell-rich regions. (D) Nestin expressing immature undifferentiated cells and proliferating immature phospho-CREB (pCREB) expressing cells were present in tumor-cell rich regions, comprising 1.8% and 0.7% of all cells, respectively. n=3. Error bars are SEM. ***p<0.001; ****p<0.0001, calculated using a Student’s t-test or Wilcoxon matched-pairs signed rank test.

## 4 Discussion

A tamoxifen-inducible Cre-recombinase under the control of the mouse Nestin upstream promoter and intron 2 enhancer was used to target two oncogenic mutations, resulting in constitutive hyperactivation of the PI3K pathway, which led to the development of brain tumors in all mice receiving tamoxifen (11). Of these mice, 12% of the male mice developed overt testicular tumors. That incidence of tumors likely reflects the incidence of nestin transgene-dependent Cre-recombinase expression. Previous studies have reported similar observations with recombination of various *loxP* alleles occurring in both germ cells and Leydig cells (19–21). Gross morphology showed testicular enlargement and regional hemorrhage, while histopathology revealed severe disruption of seminiferous tubule morphology and spermatogenesis. The features of the testicular tumors in the mice resemble human cord and stromal testicular tumors. More specifically, the testicular tumors in the mice described here, phenocopy some of the key histopathological hallmarks of human Leydig cell tumors. Malignant Leydig cell tumors in humans are typically large, exhibiting regional hemorrhage and necrosis, with malignant tissue replacing the normal testis parenchyma (22). Further, based on the cell-specificity of the *Nestin-Cre* and *Nestin-CreERT2* transgene expression, Leydig cells are one of the major extracranial target cell types (19,21,23). More recent studies propose the existence of Nestin-expressing stem Leydig cells, which represent a stem progenitor cell in the testis interstitium (24). This is a rarer testicular interstitial cell type, compared to differentiated Leydig cells, present in adult male mammals, and is more likely the cancer initiating cell in the mice.

The data presented also demonstrate that constitutive PI3K activation due to mutation of *Pik3ca* and homozygous deletion of *Pten* is sufficient to transform Leydig cells and/or stem Leydig cells into cancer cells. Although there is no reported data attributing mutations in genes associated with the regulation of the PI3K pathway in the initiation or progression of human Leydig cell tumors, mutations in two of the genes encoding the PI3K pathway p110 catalytic subunit, *PIK3CA* the *PIK3CD*, have been reported as driver genes in testicular germ cell tumors (6,25). Conditional *Pten* deletion on primordial germ cells causes testicular teratomas and *Pten* deletion, in combination with aberrant regulation of the WNT pathway drives the development of testicular granulosa cell tumors, characterized by constitutive hyperactivation of the PI3K pathway (26,27). In human Leydig cells that were chemically transformed by bisphenol A, the PI3K pathway was constitutively elevated and the cells exhibited tumor-like properties, in vitro and in vivo (28). The data presented here demonstrates that combined mutation of *Pik3ca* and homozygous *Pten* deletion are sufficient to transform stem Leydig cells into cancer initiating cells.

In humans, PTEN loss is associated with the transition from germ cell neoplasia in situ (GCNIS) to malignant germ cell tumors (29). This is consistent with our study, where *Pik3ca* mutation alone does not lead to tumor development but loss of PTEN transforms the cells and causes tumor formation. Moreover, since *PIK3CA* gene mutations are reported in 11% of KIT wild-type seminomas (6) and almost 30% of patient testicular germ cells tumors harbor somatic mutations in genes linked to the regulation of the PI3K pathway (8), the data reflect a biologically human-relevant testicular cancer model.

Immunotherapy for patients with testicular cancer remains limited, as the nature of the testicular tumor microenvironment remains poorly understood due to limited research studies conducted to date. A recent study demonstrated heterogeneous expression of the tumor cell-specific immune checkpoint biomarker, programmed death ligand-1 (PD-L1), in metastatic and post-therapy testicular cancer (30), suggesting that PD-L1 may be useful for selecting patients who may respond to immune checkpoint inhibitor immunotherapy. Our data show that in mouse testicular cancer, T-cells are the dominant tumor infiltrating immune cells, with almost no macrophage infiltration. This is similar to studies reporting that, although macrophages are the dominant immune cell type in the non-tumor adult testis, lymphocytes, including T-cells, are the dominant tumor infiltrating immune cell in testicular germ cell tumors (31). However, our data show that CD8+ T-cells comprise most (80%) of the T-cells, suggesting that therapeutic activation of cytotoxic T-cells could trigger an anti-tumor response. In testicular germ cell tumors, there are more CD4+ T-cells, compared with CD8+ T-cells (31).

Spatial analysis demonstrates that T-cells are widely distributed throughout the tumor, including tumor-cell rich regions. A minor subset of tumor cells expressed Nestin and pCREB, which is expected, since the tumor cells of origin are Nestin-expressing cells, and pCREB is a known biomarker of proliferating tumor cells (32). Moreover, Nestin expression also indicates that immature cells persist in the tumors, providing a pool of cancer stem cells which could be a cellular source for tumor recurrence. Importantly, our data demonstrates that the adaptive immune system is activated, evidenced by tumor T-cell infiltration, possibly due to Sertoli cell disruption, suggesting that our mouse model may be useful for preclinical studies testing the efficacy of immunotherapies, including chimeric antigen receptor (CAR) T-cell based therapies, which are among the most promising of the cancer immunotherapies.

## Acknowledgements

We thank members of the Mantamadiotis Laboratory for assisting in supporting various tasks associated with this study, Sarah Louise Taverner, Teresa Drever and Maya Kesar for animal care excellence, Laura Leone and the Melbourne Histology Platform and Dr Metta Jana and other members of the Centre for Advanced Histology and Microscopy, Peter MacCallum Cancer Centre for providing histopathology support.

## Author Contributions

MD prepared the manuscript draft and performed immunohistochemical analysis. LM performed multiplex immunohistochemical tissue analysis.

SSW assisted with the multiplex immunohistochemical tissue analysis, prepared figures and proofread manuscript drafts.

GF performed genotyping, immunohistochemical analysis and prepared some of the figure panels.

JA performed immunohistochemical analysis.

SD was involved in experimental design and proofreading of manuscript drafts.

SPW performed immunohistochemical analysis.

DW was involved in experimental design, provision of key materials, and proofreading of manuscript drafts.

TM funded the work, was involved in experimental design, project conceptualization, completion and submission of the manuscript.

## Ethics Statement

Mouse experiments were carried out with the approval of the Office for Research Ethics and Integrity-Animal Ethics, The University of Melbourne, School of Biomedical Sciences (AEC No 1112336.1).

## Data Availability Statement

n/a

